# Kunitz type protease inhibitor EgKI-1 from the canine tapeworm *Echinococcus granulosus* as a promising anti-cancer therapeutic

**DOI:** 10.1101/357590

**Authors:** Shiwanthi L Ranasinghe, Glen M Boyle, Katja Fischer, Jeremy Potriquet, Jason P Mulvenna, Donald P McManus

## Abstract

EgKI-1, a member of the Kunitz type protease inhibitor family, is highly expressed by the oncosphere of the canine tapeworm *Echinococcus granulosus*, the stage that is infectious to humans and ungulates, giving rise to a hydatid cyst localized to the liver and other organs. Larval protoscoleces, which develop within the hydatid cyst, have been shown to possess anti-cancer properties, although the precise molecules involved have not been identified. We show that recombinant EgKI-1 inhibits the growth and migration of a range of human cancers including breast, melanoma and cervical cancer cell lines in a dose-dependent manner *in vitro* without affecting normal cell growth. Furthermore, EgKI-1 treatment arrested the cancer cell growth by disrupting the cell cycle and induced apoptosis of cancer cells *in vitro*. An *in vivo* model of triple negative breast cancer (MDA-MB-231) in BALB/c nude mice showed significant tumor growth reduction in EgKI-1-treated mice compared with controls. These findings indicate that EgKI-1 shows promise for future development as an anti-cancer therapeutic.

## Introduction

Protein-based therapeutics enable targeted approaches for treating cancer (1). There are many benefits of proteins over small-molecule drugs mainly because of the increased surface area accessing a much wider range of protein targets (2). Protease inhibitors are important as potential cancer therapeutics as proteases are associated with carcinogenesis and cancer progression. Numerous plant protease inhibitors have recently entered human clinical trials (3). Parasites produce a range of protease inhibitors with diverse functions mainly to evade hostile adverse host reactions (4).

Several parasites, including the liver flukes, *Opisthorchis viverrini* and *Clonorchis sinensis*, and the blood fluke, *Schistosoma haematobium*, are known risk factors for cholangiocarcinoma and bladder cancer, respectively (5). In contrast, other parasites such as *Trypanosoma cruzi* (6), *Toxoplasma gondii* (7) and *Echinococcus granulosus* (8) produce metabolites with anticancer properties.

An increasing number of studies have shown that the presence of neutrophils in tumors, known as tumor associated neutrophils (TAN) correlates with poor prognosis (9), specifically in breast cancers (10). Neutrophils play major roles in tumor initiation, growth and metastasis (11) mainly through the serine protease enzyme neutrophil elastase secreted by active neutrophils. Neutrophil elastase also acts as a chemoattractant for more neutrophils (12). Therefore, potent neutrophil elastase inhibitors have stimulated much interest for development as cancer therapeutics (13).

The larval stage of the canine tapeworm (phylum Cestoda) *E. granulosus* causes echinococcosis (hydatidosis) in humans and ungulates (sheep, goats, cattle etc) when they ingest the parasite eggs containing oncospheres in contaminated food or water (14). The oncospheres hatch and penetrate the intestinal mucosa, enter the blood stream and migrate to the liver or lung. A fluid-filled larva begins to develop from a single oncosphere with subsequent formation of multiple layers, resulting in a metacestode or hydatid cyst (15). Protoscoleces, which develop asexually within the hydatid cyst, have been shown to induce the death of fibrosarcoma cells although the specific molecules involved have not been identified (8). We have shown that EgKI-1, a member of the Kunitz type protease inhibitor family, is highly expressed in oncospheres, is a potent neutrophil elastase and chemotaxis inhibitor (16) and was recently granted an International Patent Publication (17).

In this study, recombinant EgKI-1 was expressed in yeast, purified and investigated for potential anti-cancer properties *in vitro* and *in vivo*.

## Methods

### EgKI-1 expression in yeast

The pPICZαA plasmid containing the EgKI-1 gene sequence and EcoRI/ XbaI cloning sites was synthesized by Biomatik (Wilmington, USA). The plasmid (10 ng) was then transformed into *E*. *coli* XL1-Blue competent cells (Stratagene, San Diego, USA) and sequenced to confirm the integrity of the insertion. Vector bearing the confirmed sequence was inserted into *Pichia pastoris* KM71H cells using the electroporation method described by Invitrogen^TM^ (Carlsbad, USA). Briefly, a single colony of XL1-Blue cells, bearing the confirmed EgKI-1 sequence isolated from a low salt LB agar plate, was grown in 5 ml of low salt LB medium. From these cells, DNA was extracted using a Plasmid Midi kit (Qiagen, Hilden, Germany). DNA was linearized with SacI-HF (New England BioLabs, Ipswich, USA), extracted using phenol/chloroform and re-suspended in 10 mM Tris (pH 8.5) buffer. Linearized DNA (25 µg) was then mixed with 80 µl KM71H cells on ice and transferred to a 0.2 cm cuvette and an electric shock applied using Gene Pulser (Biorad, Hercules, USA). Then 1 ml of 1 M sorbitol + 200 µl HEPES mixture was added to the cells and transferred to a 10 ml tube. Cells were then incubated for 1.5 hours at 30°C, plated on YPD agar supplemented with 100 µg/ ml zeocin, and incubated at 30°C for 2-3 days. A single colony from the YPD plate was then picked and inoculated into BMGY complete medium (50 ml) and incubated at 30°C with 200 rpm agitation for 24 hours. Frozen stocks were then made with 100% sterilized glycerol and stored at -80°C for future use. The remainder of the starter culture was then used to inoculate 1 L of BMGY complete medium and grown with 200 rpm agitation for 24 hours at 30°C. On the following day, cells were harvested by centrifuging at 4000 rpm for 10 min at room temperature and re-suspended in 200 ml YNB media. Cells were grown with 200 rpm agitation for 96 hours at 30°C and then induced with 100% methanol to a final concentration of 0.5% every 24 hours to induce expression of the EgKI-1 gene under the AOX1 promoter.

### Protein purification and identification

The cultured supernatant (200 ml) was then collected by centrifugation at 10,000 rpm for 30 min at 4°C and stored at -80 ^0^C until required. The supernatant was thawed, dialyzed into 50 mM MES buffer (pH 6) and filtered through a 0.45 µm filter before being loaded on to a Hi-trap SP sepharose column (GE Healthcare Life Sciences) pre-equilibrated with 50 mM MES buffer (pH 6) (18). Unbound material was removed by washing with equilibration buffer and protein was eluted using a linear gradient of 0-1 M NaCl over 40 ml, with EgKI-1 eluting between 0.4 and 0.6 M NaCl. Purification was monitored by analysis on SDS-PAGE gels and protease inhibitory activity in fractions containing EgKI-1. Purified EgKI-1 was then dialyzed into 50 mM Tris 120 mM NaCl (pH 7) buffer and quantified using the Bradford protein assay (19). A sample of the EgKI-1 protein was visualized on 15% SDS-PAGE to verify its purity and the EgKI-1 gel band was subjected to in-gel trypsin digestion and nano high performance liquid chromatography coupled to mass spectrometry (nano LC-MS) (20).

### Cell lines

Human cancer cell lines including breast adenocarcinoma (MCF-7), breast ductal carcinoma (T47D), mammary gland epithelial adenocarcinoma (MDA-MB-231), pharynx squamous epithelial carcinoma (FaDu), cervical epithelial adenocarcinoma (HeLa), tongue squamous cell carcinoma (SCC15) and melanoma (CJM) were used to determine the effect of EgKI-1 on cell growth. Primary neonatal foreskin fibroblast (NFF) cells were used as normal human cell controls. All cells were cultured in complete media (RPMI-1640 supplemented with 10% (v/v) heat-inactivated fetal calf serum (Thermo Fisher Scientific, Waltham, USA), 3 mM 4-(2-hydroxyethyl)-1-piperazineethanesulfonic acid (HEPES)) and 100 U/ml penicillin and 100 µg/ml streptomycin (Thermo Fisher Scientific). Cell line identity was checked by Short Tandem Repeat (STR) profiling with the GenePrint^®^ 10 system (Promega, Madison, WI) according to manufacturer’s instructions. Cultured cells were routinely checked for mycoplasma infection by a specific PCR-based assay (21) and were always negative.

### Cell growth assays

Cells were seeded at 5000 cells per well in 96-well plates and incubated overnight at 37°C. On the following day cells were treated with different concentrations of the EgKI-1 protein and control wells with buffer alone and cultured for 3-5 days until the control non-treated wells were 95% confluent. Cell lines were then assayed using sulforhodamine B (SRB) to measure the inhibition of cell growth (22). The concentration of EgKI-1 needed to inhibit cell growth by 50% (IC_50_) was calculated using GraphPad Prism 7 software. The experiments were repeated three times and the mean ± SEM was determined. Growth of cancer cells was also monitored real time with the IncuCyte Zoom system in the presence of varying concentrations of EgKI-1.

### Cell migration analysis

*In vitro* scratch wound assays were carried out to determine the effect of EgKI-1 on cancer cell migration (23). This method mimics, to some extent, the migration of cells *in vivo* and is more informative and convenient than other methods(23). CJM, MDA-MB-231 and HeLa cells were grown to create a confluent monolayer in 96-well plates. Then the monolayers were scraped in a straight line to create a “scratch” using the Wound Maker-IncuCyte ZOOM-Image Lock Plate system and washed once with growth medium. After replacing the medium, cells were treated with 0.5 µM EgKI-1 or control buffer and monitored by IncuCyte Zoom. Images were acquired for each well at 12 hours interval by an in built phase contrast microscope.

The distance between one side of scratch and the other at each time point was measured using IncuCyte Zoom software. The Wound width (µm) obtained at different time points was then statistically analyzed with 2-way ANOVA using GraphPad Prism version 7.

### Immunocytochemistry

Cells were grown in 8-well tissue culture chambers on slides (Sarstedt, Nümbrecht, Germany) overnight. On the following day, different chambers were treated with EgKI-1 (2 µM) and control buffer. After 24 hours incubation, supernatants were removed and cells were fixed in ice cold methanol for 10 min followed by two washes with PBS. Cells were then permeabilized with ethanol: acetic acid (2:1) at -20°C for 5 min. After 3 × 1 min rinses in PBS, normal donkey serum (10%) was applied for 20 min at room temperature (RT). Excess normal serum was decanted and mouse anti-EgKI-1 antibody (16) and rabbit anti-tubulin antibody (Abcam, Cambridge, UK) diluted 1:80 in PBS were applied for 60 min at RT. Cells were washed with three changes of PBS and Alexa fluor donkey anti-mouse 488 and Alexa fluor donkey anti-rabbit 555 diluted 1:200 in PBS were applied for 30 min at RT. After washing with PBS, cells were stained with DAPI (1:35000 in PBS) for 2-5 min followed by another PBS wash. Prolong fluorescence mount (Thermo Fisher Scientific) was then applied to each slide and a coverslip applied. Cells were visualized under a Zeiss 780-NLO point scanning confocal microscope for the presence of EgKI-1 and tubulin. Fluorescence intensity of tubulin and total cell areas were then measured with ImageJ software (24) and statistically analyzed with GraphPad Prism version 7.

### Cell cycle phase analysis

Cells were analyzed using propidium iodide (PI) staining according to a published protocol (25). Briefly, MDA-MB-231 and HeLa cells were treated with 1 µM EgKI-1 or control buffer and cells were harvested at 24 and 48 hours after treatment. At both time points, cells in the control and EgKI-1 treated wells were collected in to 5 ml polystyrene round-bottom tubes by trypsinisation and washed thoroughly with PBS. The cells were then fixed with ice-cold 70% ethanol and stored at 4°C overnight. The ethanol was discarded after centrifugation and the cells were washed with PBS. Cells were incubated with PI/triton X-100 solution (0.02% w/v DNase free RNase, 2% w/v PI in 0.1% v/v triton X-100) for 30 min at RT and analyzed with a LSR Fortessa flow cytometer using a YG610/20 filter. ModFit LT software was used to complete the cell cycle analysis.

### Analysis of apoptosis

An Annexin V-FITC apoptosis detection kit (Biotool, York, UK) was used to determine whether EgKI-1 induced apoptosis in cancer cells. Cells were seeded at ~60% confluency in 24-well plates and EgKI-1 or control buffer treatment was carried out on the following day. After 24 hours incubation, cells were harvested by trypsinisation and washed with cold PBS. Equal amounts of Annexin V-FITC/ PI were added to the cells re-suspended in 1x binding buffer (10 mM HEPES (pH 7.4), 140 mM NaCl, 2.5 mM CaCl_2_). Cells were then incubated at room temperature for 15 min and analyzed with a BD LSR Fortessa flow cytometer using a YG610/20 filter for PI and a B530/30 filter for FITC. FlowJo version 10 was used to analyze cell staining and to determine the percentage positivity. Annexin V^+^/PI^-^ cells were considered as early apoptotic cells and Annexin V^+^/PI^+^ cells were considered as late apoptotic cells.

### Proteomics quantitative analysis following EgKI-1 exposure

To determine which proteins in MDA-MB-231 cancer cells were affected by EgKI-1 treatment, quantitative SWATH-MS (sequential window acquisition of all theoretical spectra-mass spectrometry) was used as described by Gill *et al*(26) was used with few modifications. Cells were harvested prior to treatment as the controls, and after 0.5, 2, 4, 8 and 24 hours of EgKI-1 treatment. Protein extraction and tryptic peptide fragment generation was conducted following a modified FASP protocol for high-throughput sample preparation (27). The data for the ion library generation were obtained using Data Dependent Acquisition (DDA) on each individual samples as previously described (28) while quantitative data from different time points was obtained using SWATH acquisition with the same conditions used in the DDA experiments. A rolling collision energy method was used to fragment all ions in a set of 26 sequential overlapping windows of 25 AMU over a mass range coverage of 350-1,000 (m/z). Data was acquired and processed using Analyst TF 1.7 software (AB SCIEX).

### Protein library generation and Bioinformatic analysis of SWATH protein quantification

Spectral searches of processed LC-MS/MS data were performed using ProteinPilot v4.5 (AB SCIEX) using the Paragon algorithm (version 4.5.0.0). Background correction was used and biological modifications specified as an ID focus. The detected protein threshold was set as 0.5 and the false-discovery rate (FDR) was calculated using searches against a decoy database comprised of reversed sequences. Searches were conducted against the UniProt human reference proteome set comprising 70953 protein sequences.

For spectral library generation and SWATH XIC peak area extraction PeakView v2.2.0 (AB SCIEX) with the SWATH acquisition MicroApp was used with ion library parameters set to 6 peptides per protein, 6 transitions per peptides, a peptide confidence threshold of 99% and FDR threshold to 1%. The XIC time window was set to 6 min and XIC width to 75 ppm. All SWATH experiments used iRT calibrants to normalize retention times. To generate the quantitation table files for ions, peptides and proteins, Marker View v1.2.1.1 (AB SCIEX) was used and the relative area under peaks across the different experiments was normalized based on the iRT internal calibrant.

### Bioavailability of EgKI-1

To determine the absorption rate and stability of EgKI-1 in serum, 50 µg of the protein in 50 µl of Tris/NaCl buffer (pH 7) was injected intraperitoneally into BALB/c mice and blood samples were collected at 5, 30, 120, and 300 min after injection. The serum was separated and samples analyzed using liquid chromatography-mass spectrometry to monitor the presence of EgKI-1 at the different time points.

### *In vivo* animal model

MDA-MB-231 cells (1×10^6^) were injected into the right inguinal mammary tissue of 6-7 weeks old BALB/c nude mice. When the tumors reached approximately 30 mm^3^, in 20 days, mice in the control group received buffer only (150 mM NaCl, 20 mM Tris HCl, pH 7) and the treatment group were treated with 4 mg/kg of EgKI-1 per mouse (equivalent to 80 µg per 20 g mouse). EgKI-1 and buffer were injected into the tumors in a total volume of 50 µl once a day every other day. All mice were monitored daily and tumor volumes were measured twice a week using digital Vernier caliper and expressed as mm^3^ according to the formula, a × b × b × 0.5, where “a” the length and “b” the measured breadth of the tumor. Mice were also assessed for clinical signs according to an approved clinical score sheet for distress during the course of the experiment (29). Scores for each parameter were summed to give a possible total of 8. Less than 3 was considered a mild clinical score, between 3-6 a moderate and over 6 was considered a severe clinical score. Experimentation on an individual mouse was terminated when an unacceptable clinical score (>6) was reached, or the cumulative tumor burden of the animal exceeded 1000 mm^3^. Mice were humanely euthanized by exposure to carbon dioxide at the end of the experiment and tumors were collected.

Paraffin blocks were made with the harvested tumor tissues and sections were stained with mouse monoclonal Ki67 antibodies for immunolabelling screening in the QIMRB histology facility. Ki67 is a nuclear protein that is expressed in proliferating cells and is thus a marker for cell proliferation in solid tumors (30). Ki67 stained slides were then scanned at 20× magnification with an Aperio Scanscope XT slidescanner and digital images were analyzed with ImageScope viewing software. The Aperio nuclear algorithm which is based on the spectral differentiation between brown (positive) and blue counter staining was used for analysis. Total percentage positivity for each slide was then calculated and analyzed using GraphPad Prism.

### Animal ethics statement

This study was performed in strict accordance with protocols approved by the QIMRB Animal Ethics Committee, approval number A1606-617M, which adheres to the Australian code of practice for the care and use of animals for scientific purposes, as well as the Queensland Animal Care and Protection Act 2001; Queensland Animal Care and Protection Regulation 2002. Mice were housed in a specific pathogen free facility with 12 hours light/dark cycle and continual access to food and water.

### Statistical analysis

All data are presented as the means ± standard mean of error (SEM) of three different experiments. A P-value of <0.05 was considered as statistically significant according to the Student’s t-test and ANOVA tests. All statistical analysis was performed using GraphPad Prism version 7.

### Data availability

All data generated or analyzed during this study are included in this published article (and its Supplementary Information files).

## Results

### Recombinant EgKI-1 expression and purification

In order to examine the effects of EgKI-1 on cancer cell lines the protein was expressed in yeast. The denaturing SDS-PAGE of the purified EgKI-1 protein revealed a single band around 8 kDa which is consistent with the predicted molecular size of 8.08 kDa (S1 Fig). MALDI-TOF MS analysis with the final purified protein sample showed a 100% intensity peak at 7.506 kDa. In-gel digestion identified the protein band as a BPTI/Kunitz inhibitor from *E*. *granulosus* with 91% coverage.

### Recombinant EgKI-1 inhibits cell growth *in vitro*

To test the effects of EgKI-1 on the proliferation of cancer cell lines, end point and real time monitored cell growth assays were carried out. EgKI-1 treatment inhibited the growth of a range of human cancer cells at different rates when assessed by end point SRB assay (S2 Fig; Table 1). The protein had substantially less effect on the growth of primary neonatal foreskin fibroblast (NFF) cells, and had ~9-40 fold high IC_50_ value compared with the IC_50_ values of tested human cancer cell lines.

**Table 1:** IC_50_ values for different human cancer cell lines tested with EgKI-1.

| IC<sub>50</sub> values (μM) with 95% CI | | | | | | | | |
| --- | --- | --- | --- | --- | --- | --- | --- | --- |
| Cell line | NFF | MCF-7 | MDA-MB-231 | CJM | SCC15 | FaDu | HeLa | T47D |
|  | 47.134 | 2.075 | 5.162 | 4.660 | 4.025 | 2.674 | 1.157 | 3.012 |
|  | 18.2 to | 1.88 to | 4.16 to 6.10 | 3.12 to | 3.69 to | 2.17 to | 0.9 to | 2.64 to |
|  | 59.4 | 2.27 |  | 8.33 | 4.41 | 3.28 | 1.49 | 3.36 |

CI - Confidence Interval

Real time monitoring using the IncuCyte showed that cells treated with 1 µM EgKI-1 eventually lost their typical shape, lost their ability to proliferate with time (Fig 1a), and exhibited dose-dependent growth inhibition compared with the control cells (Fig 1b). Furthermore, the IncuCyte analysis showed that cells continued to grow in the presence of EgKI-1, at a concentration of around 0.5 µM, before starting to die after ~60 hours making it a more sensitive assay than end point staining.

**Fig 1:**
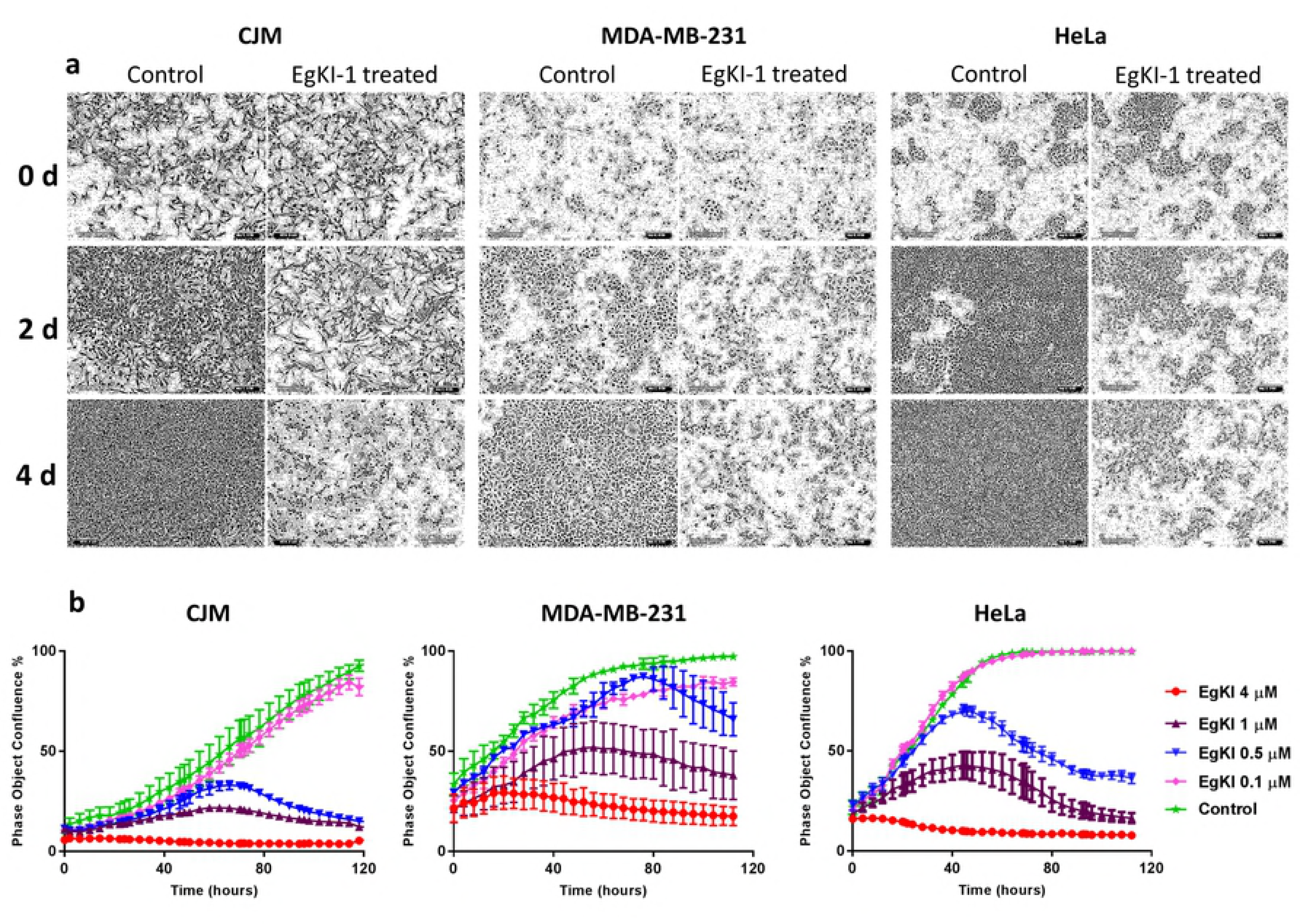
Growth of cancer cells inhibited by EgKI-1. **(a)** CJM, MDA-MB-231 and HeLa cells lost shape and started dying when treated with 1 µM EgKI-1 (0.8 µg in 100 µl/ well) over time. **(b)** Dose-dependent growth inhibition curves with varying concentrations of EgKI-1.

### EgKI-1 treatment inhibits cell migration

We examined the effect of EgKI-1 treatment on cell migration on a panel of representative cell lines. The real time scratch wound assay analysis using the IncuCyte system revealed that 0.5 µM EgKI-1 inhibited the migration of the CJM (melanoma), MDA-MB-231 (breast cancer) and HeLa (cervical cancer) cells *in vitro* (Fig 2a). The gap in the scratch wound in control wells decreased eventually with time. The wound width was significantly larger in EgKI-1-treated wells compared with controls after 2 days indicating the inhibition of cell migration by EgKI-1. According to the analysis, the closure of the scratch in EgKI-1-treated cells were 12% in CJM, 18% in MDA-MB-231 and 16% in HeLa compared with 60%, 58% and 100% in the controls respectively (Fig 2b). This data indicates that EgKI-1 inhibits cancer cell migration.

**Fig 2:**
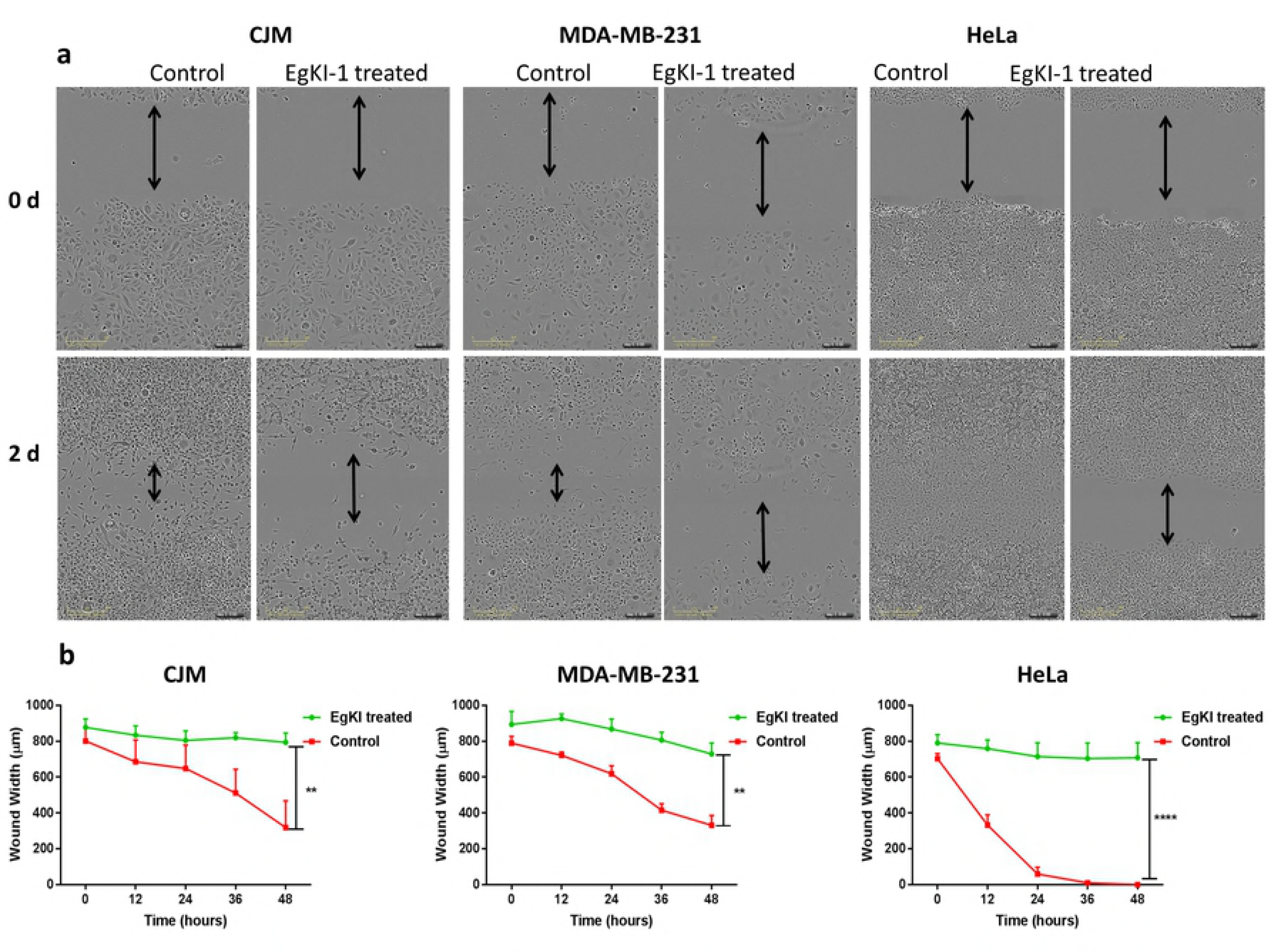
EgKI-1 inhibits cancer cell migration *in vitro*. In the control wells the gap closed with time but in the treated wells the gap closure was significantly reduced compared with the controls. P<0.01** and p<0.0001****

### Immunocytochemistry demonstrates EgKI-1 internalization

The presence of green fluorescence in the cells indicated that cancer cells internalized the EgKI-1 protein after treatment. Furthermore, vacuoles started to appear in the cytoplasm, which can lead to cell membrane collapse and eventually necrosis. There were significant decreases in the intensity of tubulin and total cell area indicating the degradation of the cytoskeleton following treatment with the EgKI-1 protein (Fig 3).

**Fig 3:**
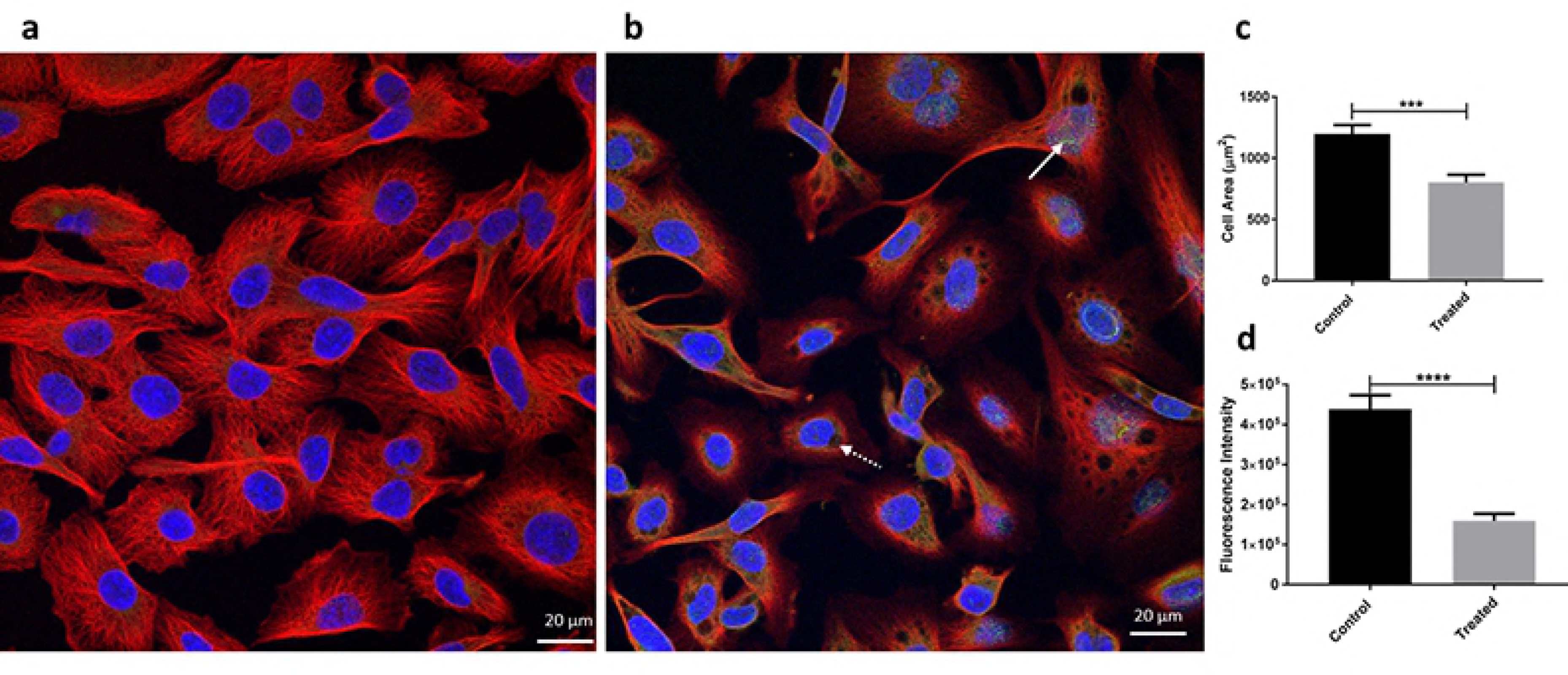
Immunocytochemistry. **(a)** Control cells; **(b)** EgKI-1 treated MDA-MB-231 breast cancer cells 24 hours post-treatment. White arrow shows the green in treated cells indicating the EgKI-1 protein; White dotted arrow indicates the vacuole in the cell; Red indicates tubulin in cytoplasm; Blue, DAPI nuclear stain; **(c)** Cell area reduction (p<0.001***) and **(d)** Red fluorescence intensity of tubulin (p<0.0001****) after analysis by Student’s t-test.

### EgKI-1 disrupts cell cycle profile

Cell cycle profiles of MDA-MB-231 and HeLa cells were assessed after EgKI-1 treatment. Even though significant non-adherent cells were observed only attached cells were harvested and assessed. EgKI-1 treatment caused a significant increase in S phase in both MDA-MB-231 and HeLa cell lines 2 days post-treatment. MDA-MB-231 cells treated with EgKI-1 (2 µM) significantly decreased the G2/M phase (~60%) by arresting the proliferative S phase of the cell cycle at 2 days post-treatment compared with the control cells (Fig 4). In HeLa cells, EgKI-1 perturbed cell cycle progression by significantly decreasing the non-proliferative G0/G1 fraction (~30%) and increasing the S (~30%) and G2/M (~80%) fractions 2 days post-treatment (Fig 4). No significant difference was observed 1 day post-treatment (S3 Fig).

**Fig 4:**
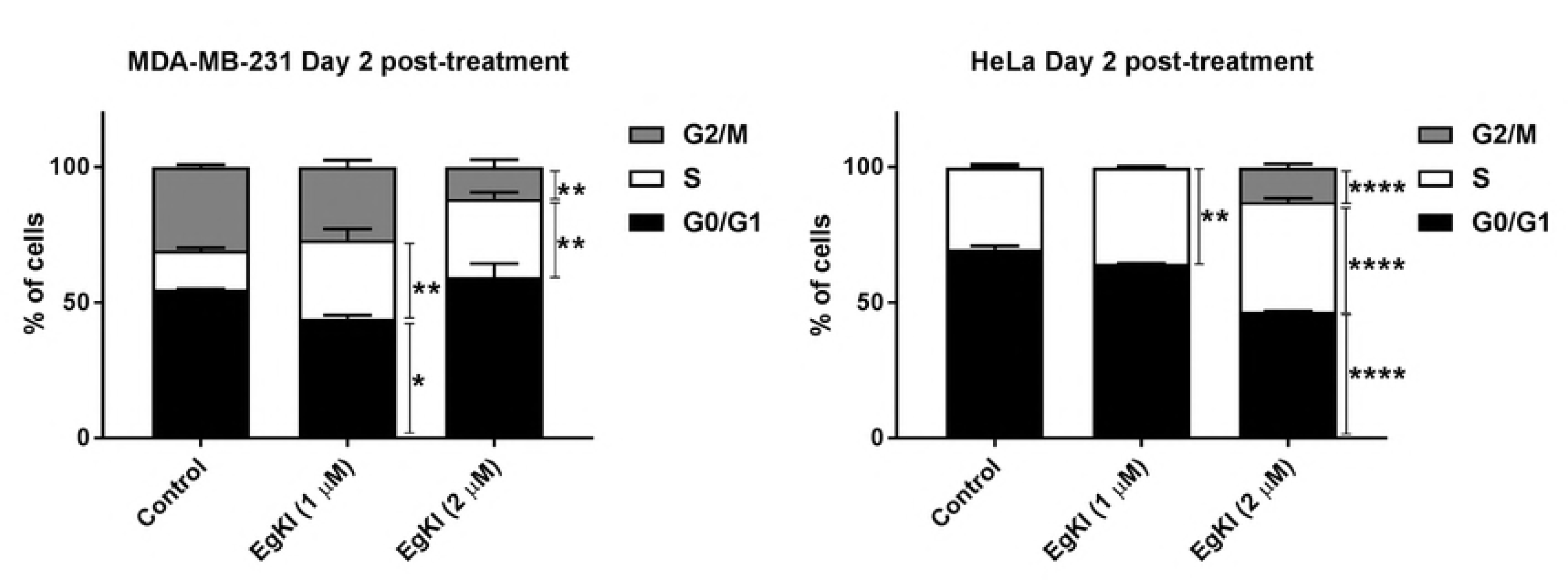
EgKI-1 induces cell cycle arrest. MDA-MB-231 and HeLa cells were treated with 1 µM and 2 µM EgKI-1 for 48 hours and the cell cycle distribution was determined by flow cytometry. Results represent the mean ± SEM from 3 independent experiments. p<0.05 *, p<0.01 **, p<0.001 *** and p<0.0001**** by 2 way ANOVA test.

### EgKI-1 can induce apoptosis

As we observed a significant proportion of non-adherent cells following EgKI-1 treatment, treated cells were assessed for apoptosis. EgKI-1 induced apoptosis in MDA-MB-231 breast cancer cells 24 hours post-treatment in a dose-dependent manner. Cells treated with 0.2 µg/well of EgKI-1 (2 µM) had a significantly higher percentage of cells (19.9%) in early apoptosis compared with control cells whereas cells treated with 0.8 µg/well EgKI-1 (0.5 µM concentration) had a higher percentage of cells (78.6%) in late apoptosis (Fig 5).

**Fig 5:**
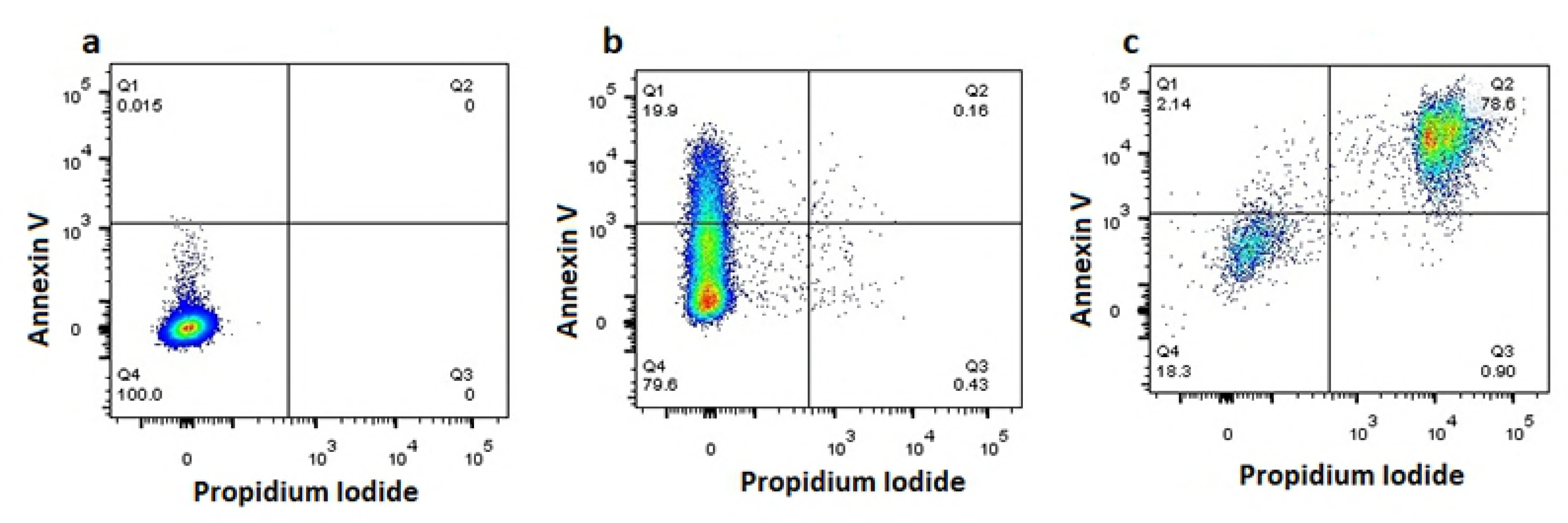
Induction of apoptosis in MDA-MB-231 breast cancer cells following EgKI-1 treatment. **(a)** Control, untreated cells were alive, being Annexin V**^-^**/PI**^-^**; **(b)** 20% of cells treated with the lower EgKI-1 concentration (0.25 µM) were in early apoptosis (Annexin V^+^/PI**^-^**); **(c)** Cells treated with the higher EgKI-1 (1 µM) concentration exhibited the highest percentage of late apoptotic/ early necrotic cells (Annexin V^+^/PI^+^).

### Proteomics analysis following EgKI-1 treatment

SWATH-MS analysis identified a total of 1770 proteins (shown in the heat map; S4 Fig) in cancer cells. EgKI-1 treatment mainly up-regulated Bcl-2-like protein 13 (Q9BXK5) and inhibitor of nuclear factor kappa-B kinase subunit beta (O14920) expression with time. Tetraspanin (H7BXY6), nucleoside diphosphate kinase A (P15531) and double-strand-break repair protein (O60216) expression were down-regulated with time (S5 Fig).

### Bioavailability of EgKI-1

When injected intraperitoneally into mice, the EgKI-1 protein was absorbed into the blood in 5 minutes and the highest serum availability occurred within 30 minutes. Most of the EgKI-1 protein had been cleared from the blood by 5 hours (S6 Fig). No adverse or toxic reactions resulting from the administration of EgKI-1 to the mice occurred.

### EgKI-1 reduces tumor growth in an in vivo breast cancer model

We assessed the impact of EgKI-1 intra-lesional treatment on the growth of MDA-MB-231 tumors *in vivo*. The tumors were allowed to reach approximately 30 mm^3^ before treatment with 4 mg/kg of EgKI-1, or equivalent vehicle, once per day every other day. The growth of MDA-MB-231 breast tumors was significantly reduced by 57.7% in EgKI-1-treated mice compared with control mice (Fig 6). Furthermore, expression of Ki67 protein, which is a proliferation marker, was significantly reduced in the EgKI-1-treated tumor tissues indicating ~80% reduction of tumor proliferation (Fig 6) as suggested by the *in vitro* data.

**Fig 6:**
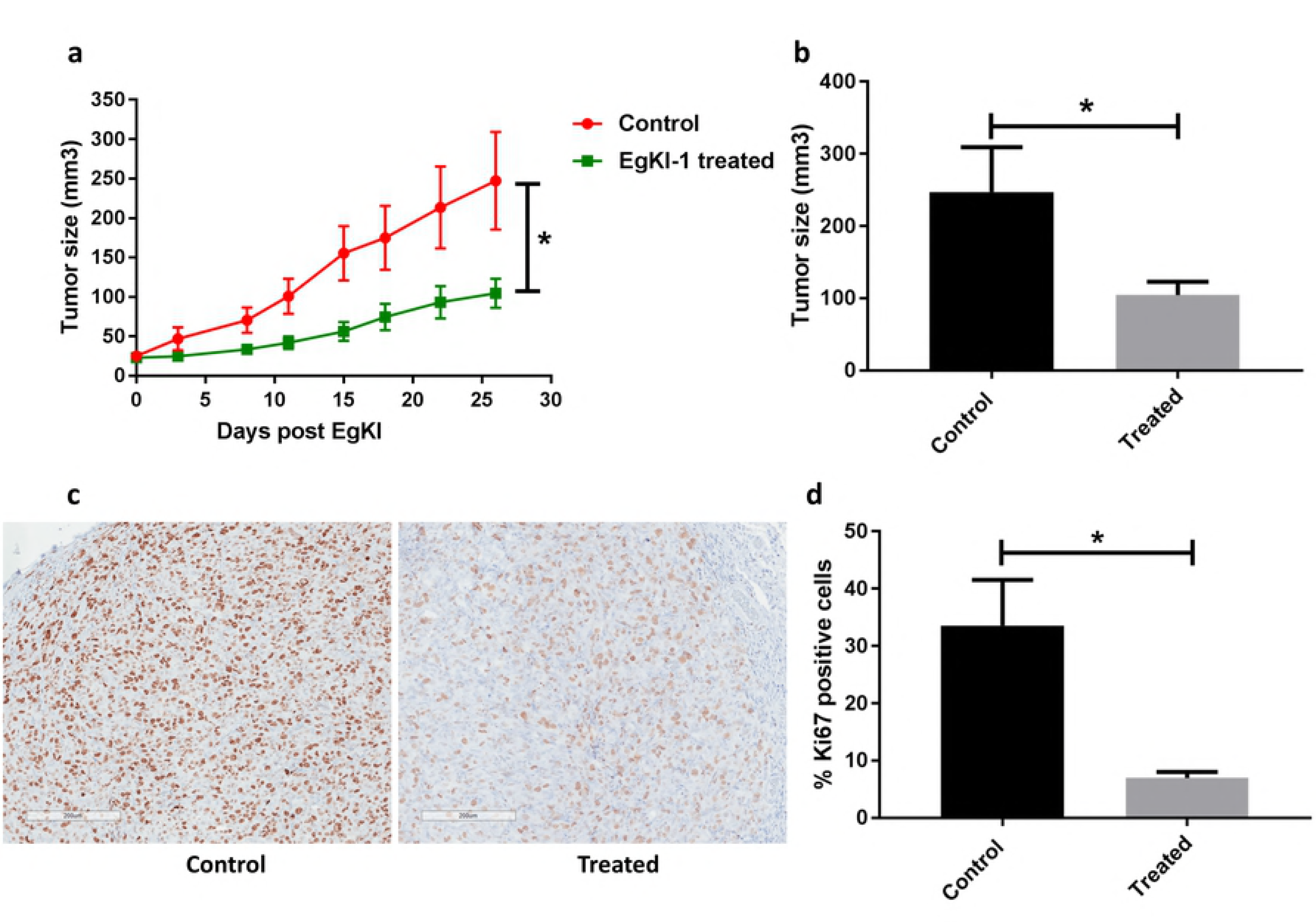
Anti-tumor effects of EgKI-1 treatment *in vivo*. **(a)** Tumor growth curve of MDAMB-231 tumor bearing mice receiving EgKI-1 treatment (4 mg/kg) compared with control mice (P<0.05* by Student’s t-test). **(b)** Tumor size of EgKI-1 treated (n=9) and control mice (n=6) surviving at the end of the experiment (P<0.05* by Student’s t-test) showing ~60% reduction of tumor growth in treated mice. **(c)** Representative bright field images for the immunohistochemical staining of Ki67 proliferation marker (brown) in control and treated tumors (scale bar indicates 200 µm). **(d)** Quantification of Ki67-positive cells (P<0.05* by Student’s t-test).

## Discussion

This study describes the broad-spectrum anti-cancer activities of EgKI-1 which is the first anti-cancer molecule identified from the cestode *E*. *granulosus*. Although we hypothesized based on our previous studies (16) that the potent neutrophil elastase activity of EgKI-1 would result in cancer cell regression, the *in vitro* culture medium we employed did not contain neutrophils or neutrophil elastase (S2 Fig). Therefore there must be additional mechanisms whereby EgKI-1 adversely affects cancer cell growth. Inhibition of the growth of the different cancer cell lines with an IC_50_ range of 1.1 - 5.1 µM, with only minimal reduction in cell growth of primary neonatal fibroblasts, suggests that EgKI-1 potentially interacts with cancer-specific proteins only. EgKI-1 treatment specifically inhibited the growth of invasive, rapid proliferative cell lines like MCF-7, FaDu and HeLa with lower IC_50_ values compared with other tested cell lines (31, 32). Cancer cell migration can lead to tumor metastasis and invasion (33). Therefore the observed inhibition of cancer cell migration by EgKI-1 is also an important aspect to consider for cancer therapeutic development.

Further, the apparent cell growth inhibition by EgKI-1 was due to an increase in apoptosis, potentially by disrupting cell cycle progression. Both HeLa and MDA-MB-231 exhibited increased numbers of cells in S phase of the cell cycle following treatment. While MDA-MB-231 showed a reduction of cell numbers in G2/M transition, HeLa had increased G2/M numbers following treatment. Abnormal, deregulated cancer cells undergo unrestricted division which is different from other cancers and, as an “immortal” cell line, HeLa cells divide differently (34). This can possibly be a reason for the changes in the cell cycle of HeLa and MDA-MB-231 cell lines treated with EgKI-1. Identifying new cancer-specific molecules that can target mitosis can optimize combinatorial treatments in the future (35). The induction of apoptosis by EgKI-1 at the same concentration that led to 50% inhibition in cell growth/proliferation shows the effectiveness of the protein in cancer cell growth inhibition. Apoptosis-targeted cancer therapy has undoubtedly been an indispensable approach in treating cancer, in order to make damaged cells commit suicide (36). The immunocytochemistry results presented here indicated that EgKI-1 was internalized in cancer cells. The immediate effects on the plasma membrane evident after EgKI-1 treatment might therefore support the sequence of events involving EgKI-1 in the mechanism of action leading to the induction of apoptosis.

SWATH-MS analysis supports the relative quantification of large fractions of a proteome in a single sample (37). EgKI-1 treatment up-regulated BcL-2 like protein 13 (UniProt IDQ9BXK5) which promotes the activation of caspase-3 and apoptosis (38). EgKI-1 treatment further down-regulated the expression of tetraspanin (UniProt ID-H7BXY6) and BRCA1-associated ATM activator-1 (BRAT1) (UniProt ID-Q6PJG6) proteins. Tetraspanins can promote multiple cancer stages by playing key roles in tumor initiation and metastasis (39). BRAT1 plays broad roles in DNA repair and cell cycle regulation involved in controlling cell growth (40). Therefore, down-regulation of tetraspanin and BRAT1, caused by EgKI-1, can contribute to cancer cell growth inhibition but additional study is required to identify the precise molecules and mechanisms of action of EgKI-1 in cancer cell growth inhibition.

Metastatic breast cancer is the leading cause of cancer death in women worldwide (41). Therefore, the MDA-MB-231 cell line was selected for further investigation and in *in vivo* animal model studies. In this study we used intra-tumor delivery of EgKI-1 as direct injection into a tumor lesion has the advantage that much higher drug concentrations can be applied at the tumor site with the result that intra-lesional anti-tumor therapeutics are considered as more effective than other routes (42). After 26 days of EgKI-1 treatment there was ~60% reduction of tumor growth in treated mice compared with controls. As Ki67 expression is strongly associated with tumor cell proliferation (30), the significantly decreased numbers of Ki67-positive cells in tumor tissues treated with EgKI-1 reflect its inhibitory effect on cell proliferation *in vivo*.

A therapeutic with a molecular size less than 10 kDa allows rapid extravasation from blood vessels and rapid transport to tumor targets resulting in maximal tumor uptake (43). However, such molecules have shorter serum half-lives than longer peptides as they are quickly cleared by renal filtration (44). PEGylation, the technique of covalently attaching polyethylene glycol (PEG) to a molecule, which may be a low molecular size protein, enzyme or nanoparticle, has proven to be one of the best methods for the passive targeting of anti-cancer therapeutics (45, 46). Accordingly, this technology will be applied in the future to improve the serum half-life of EgKI-1 if needed.

Another future consideration will be to monitor the up or down regulation of genes involved in cancer progression and metastasis following EgKI-1 treatment. Further, EgKI-1 treatment can inhibit the cellular activity of different proteases, including trypsin, plasma kallikrein and cathepsins, and matrix metalloproteases as analyzed with P237, P139 and P126 fluorescence substrates (unpublished data). We have also shown that EgKI-1 can induce some anti-inflammatory cytokines (unpublished data), which is important as chronic inflammation increases the risk of cancer and a reduction in inflammation helps in cancer therapy (47). Therefore immunomodulation effects induced by EgKI-1 may also play a role in cancer growth inhibition *in vivo*, an area that needs to be explored in the future using immunocompetent mouse models of cancer.

## Conclusion

The selective killing of malignant cells without affecting normal cells is the ultimate goal of cancer therapy. As a low molecular weight (<10 kDa) protein, EgKI-1 has potential to penetrate tumor tissues for effective interaction and killing of malignant cells. EgKI-1 treatment significantly reduced the rate of breast cancer growth *in vivo* and represents a promising molecule for development as a future anti-cancer therapeutic.

## Author Contributions Statement

SLR, GMB, JPM and DPM planned the study. SLR, KF and JP performed the experiments, analyzed the data and prepared figures. GMB and DPM provided guidance to the study and corrected the manuscript. All authors reviewed the manuscript.

## Supporting Information

**S1 Fig: Purified recombinant EgKI-1 on an SDS-PAGE gel.** 1, 3 µg EgKI-1; L, molecular size markers; 2, 6 µg EgKI-1.

**S2 Fig: Viability of different cancer cell lines in the presence of EgKI-1.** Viable cell percentages of primary neonatal foreskin fibroblast cells (NFF), breast cancer cell lines (MCF-7, MDA-MB-231, T47D), melanoma (CJM), squamous cell carcinomas (SCC15, FaDu) and cervical adenocarcinoma (HeLa) cell lines with different EgKI-1 concentrations *in vitro*.

**S3 Fig: EgKI-1 effect on cell cycle arrest.** MDA-MB-231 and HeLa cells were treated with 1 µM and 2 µM EgKI-1 for 24 hours and the cell cycle distribution was determined by flow cytometry. No significant difference was observed among the different cell cycle phases.

**S4 Fig: Differentially expressed proteins quantified by SWATH-MS analysis.** Heat map analysis of 1770 proteins among three biological replicates of the control and EgKI-1 treated samples after 30 min, 2 hours, 4 hours and 24 hours. The fold change value of the MS signal intensity is shown.

**S5 Fig: Fold change of protein expression which down/up regulated with time for 24 hours.** H7BXY6:Tetraspanin, P15531:nucleoside diphosphate kinase A, O60216:double-strand-break repair protein, O14920:inhibitor of nuclear factor kappa-B kinase subunit beta, Q9BXK5:Bcl-2-like protein 13.

**S6 Fig: Serum stability of EgKI-1.** After intraperitoneal injection into mice, EgKI-1 was absorbed into the blood within 5 minutes and had been cleared in 5 hours.

